# GiniClust3: a fast and memory-efficient tool for rare cell type identification

**DOI:** 10.1101/788554

**Authors:** Rui Dong, Guo-Cheng Yuan

## Abstract

**Motivation:** With the rapid development of single-cell RNA sequencing technology, it is possible to dissect cell-type composition at high resolution. A number of methods have been developed with the purpose to identify rare cell types. However, existing methods are still not scalable to large datasets, limiting their utility. To overcome this limitation, we present a new software package, called GiniClust3, which is an extension of GiniClust2 and significantly faster and memory-efficient than previous versions.

**Results:** Using GiniClust3, it only takes about 7 hours to identify both common and rare cell clusters from a dataset that contains more than one million cells. Cell type mapping and perturbation analyses show that GiniClust3 could robustly identify cell clusters.

**Availability:** GiniCluster3 is implemented in the open-source python package, with source code freely available through the Github (https://github.com/rdong08/GiniClust3).

**Supplementary information:** Supplementary data are available at Bioinformatics online.

## 1 Introduction

The rapid development of single cell technologies has greatly enabled biologists to systematically characterize cellular heterogeneity (see reviews (Kolodziejczyk, et al., 2015; Papalexi and Satija, 2018; Stegle, et al., 2015; Yuan, et al., 2017)). While many methods have been developed to identify cell types from single cell transcriptomic data (Butler, et al., 2018; Wolf, et al., 2018; Zheng, et al., 2017), most are designed to identify common cell types. As the throughput becomes much higher, it is also of considerable interest to specifically identify rare cell types. Several methods have been developed (Grun, et al., 2015; Grun, et al., 2016; Jiang, et al., 2016; Jindal, et al., 2018; Tsoucas and Yuan, 2018); however, existing methods are not scalable to very large datasets. Considering the fact that atlas-scale datasets may contain hundreds of thousands or even millions of cells (Han, et al., 2018; Rozenblatt-Rosen, et al., 2017; Zeisel, et al., 2018; Zheng, et al., 2017), there is an urgent need to develop faster method for rare cell type detection.

In previous work, we developed GiniClust to identify rare cell clusters, using a Gini-index based approach to select rare cell-type associated genes (Jiang, et al., 2016). Recently, we extended the method to identify both common and rare cell clusters, using a cluster-aware, weighted ensemble clustering approach (Tsoucas and Yuan, 2018). These methods have been used to analyze datasets containing up to 68,000 cells. Here we have further optimized the algorithm so that it can be efficiently used to analyze dataset containing over one million cells. By using a real single-cell RNA-seq dataset as an example, we show that this new extension, which we call GiniClust3, can efficiently and accurately identify both common and rare cell types.

## 2 Methods

### 2.1 Data source

A mouse brain single-cell RNA-seq dataset was downloaded from 10X genomics website: (https://support.10xgenomics.com/single-cell-gene-expression/datasets/1.3.0/1M_neurons). This dataset contains 1.3 million cells obtained from cortex, hippocampus and ventricular zones of E18 mice. Raw data was pre-processed by using Scrublet (Wolock, et al., 2019) (version 0.2.1) to remove doublets with default setting. The resulting data was further filtered to remove genes expressed in fewer than ten cells and cells expressed fewer than 500 genes. Raw UMI counts were normalized by Scanpy (Wolf, et al., 2018) with the following parameter setting: sc.pp.normalize_per_cell (counts_per_cell_after=1e4). A total number of 1,244,774 cells and 21,493 genes passed this filter were retained for further analysis.

### 2.2 Details of GiniClust3 pipeline

The overall strategy is similar to GiniClust2 (Tsoucas and Yuan, 2018). The implementation of each step is optimized to improve computation and memory efficiency (Fig. 1a). The details of the GiniClust3 pipeline are as follows.

**Fig. 1:**
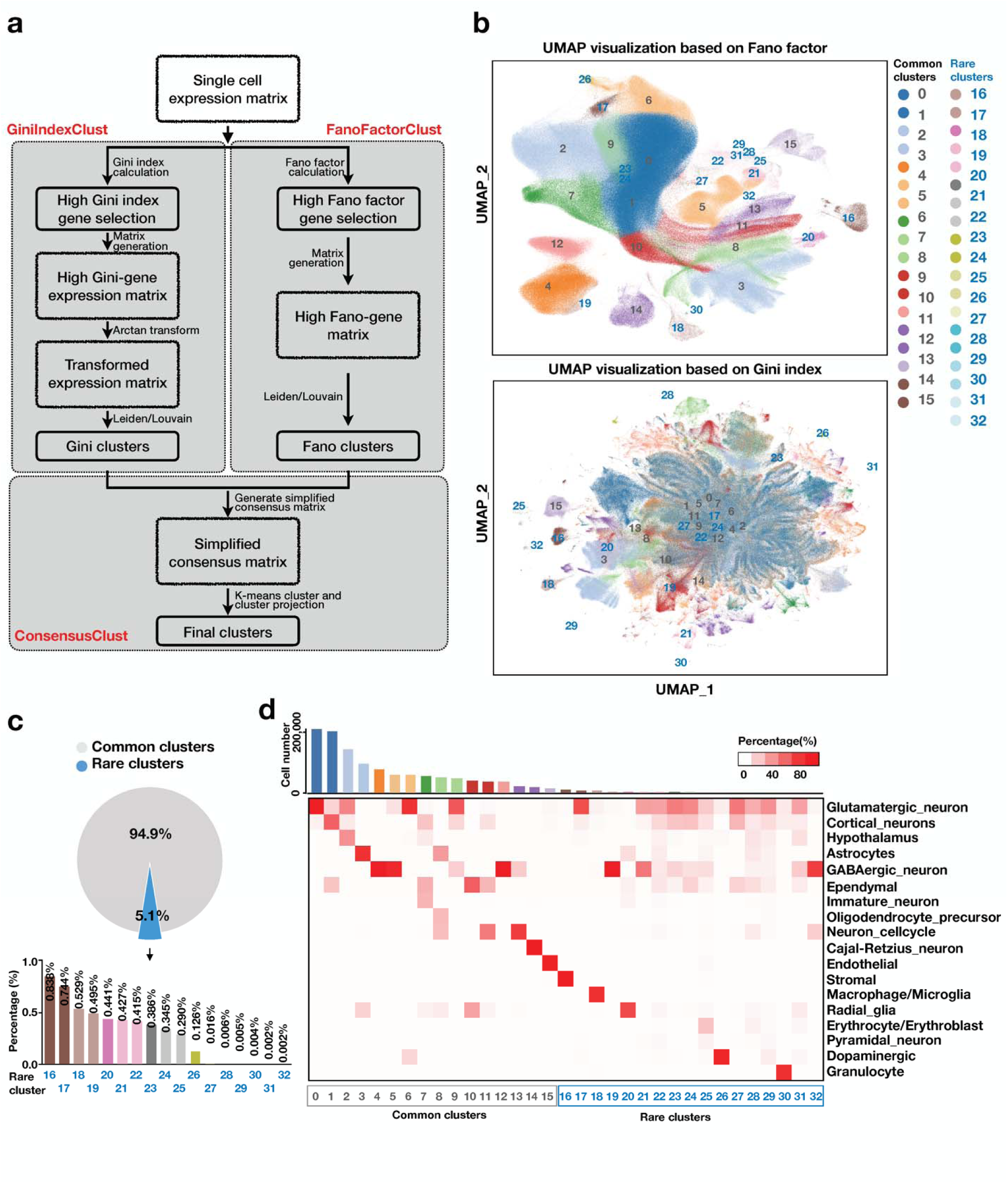
(a) An overview of the GiniClust3 pipeline. Input single-cell expression matrix is clustered based on features selected by Gini index (GiniIndexClust) and by Fano factor (FanoFactorClust), respectively. The results are then integrated using a cluster-aware, weighted consensus clustering algorithm (ConsensusClust). (b) UMAP visualization of the gene expression patterns based on Fano-factor (top) and Gini index (bottom) selected features, respectively. Consensus clustering results are indicated by different colors. (c) The proportion of rare cell cluster in entire population. (d) Heatmap of cell type mapping of common and rare clusters from scMCA analysis. Bar plot in the top indicates the cell number for each cluster.

#### Step 1: Clustering cells using Gini index-based features

a. *Gini index calculation and normalization*. After data pre-processing, the Gini index for each gene is calculated as twice of the area between the diagonal and Lorenz curve, as described before (Jiang, et al., 2016). The range of Gini index values is between 0 to 1. Then, Gini index values are normalized by using a two-step LOESS regression procedure as described before. Genes with Gini index value ≥ 0.6 and *p* value < 0.0001 are labeled as high Gini genes and selected for further analysis.
b. *Cell cluster identification by Leiden algorithm*. In previous versions (Jiang, et al., 2016; Tsoucas and Yuan, 2018), DBSCAN was used for clustering. While DBSCAN is effective for identify rare cell clusters, this method is both time and memory consuming. In GiniClust3, we replace DBSCAN with the Leiden clustering algorithm (Traag, et al., 2019), which is known for improved numerical efficiency. Alternatively, users can also select the Louvain clustering algorithm (Blondel, et al., 2008) by setting “method=louvain”. The neighbor size we set in Gini index-based clustering of mouse brain single-cell dataset is 15 (n_neighbors=15). Lower threshold for neighbor size to efficiently identify rare clusters in smaller datasets is recommended.

#### Step 2: Clustering cells using Fano factor-based features

Highly variable genes are identified by using Scanpy. These genes are used to identify common cell clusters by using principal component analysis (PCA) followed by Leiden or Louvain clustering, using the default settings in Scanpy (Wolf, et al., 2018). The neighbor size we set in Fano factor-based clustering of mouse brain single-cell dataset is 15 (n_neighbors=15).

#### Step 3: Combining the clusters from Steps 1 and 2 via a cluster-aware, weighted consensus clustering approach effectively

The weighted consensus clustering method is described before (Tsoucas and Yuan, 2018). In brief, connectivity of cells in different cluster results (*P*^*G*^ and *P*^*F*^) are calculated. We set the cell-specific weights for the Fano factor-based clusters *w*^*F*^ as a constant value *f’* while the cell-specific GiniIndexClust weight *w*^*G*^ are determined as a logistic function of the size of cluster containing the particular cell. After normalization of the *w*^*F*^ and *w*^*G*^, we calculated the consensus value based on the weight and connection.

To improve computational efficiency, we introduce an alternative, but mathematically equivalent implementation. This is based on a simple observation that the consensus matrix has a high degree of redundancy: many entries have exactly the same values because they correspond to cells from same Gini and Fano cluster annotations. To remove this redundancy, we create an equivalent consensus matrix where each entry corresponds to a combination of clustering annotations. This new consensus matrix is much smaller than the original one, therefore increasing computational efficiency. K-means clustering is applied to the new consensus matrix, then the results are easily converted back to single-cell level clustering. Finally, clusters with cell population <1% are considered as rare clusters.

## 3 Results

To test the utility of GiniClust3, we applied the method to analyze a public single-cell RNA-seq dataset containing 1.3 million single cells obtained from three regions in the mouse brain (see Methods for details). After filtering out lowly-expressed genes and poor-quality cells (such as those likely to be doublets), a 1,244,774 cell-by-21,494 gene count matrix was left for further analysis. We next sought to characterize the identities of cell populations by using GiniClust3. A total number of 16 common and 17 rare cell clusters (cell population < 1%) were identified (Fig. 1b, s1a), with the smallest cluster containing only 21 cells (cell population = 0.002%) (Fig. 1c and Table s1). The total time of cluster identification for both common and rare cell took ~7-hour time, and 103G memory on a Xeon E5-2683 with 56 threads and 640GB memory server, indicating GiniClust3 is suitable for analyzing very large datasets.

To annotate these cell clusters, we mapped each cluster to mouse cell atlas (MCA) (Han, et al., 2018) by using the scMCA algorithm (Sun, et al., 2019). Ten of the sixteen common clusters (cluster 0, 1, 4, 5, 6, 9, 12, 13, 14 and 15) were mapped to specific cell types in MCA with expected abundance. These include glutamatergic neurons, astrocytes, GABAergic neuron, ependymal, cell cycle neuron, cajal-retzius neuron and endothelial (Fig. 1d). For example, cluster 0 is mapped to glutamatergic neurons, which are known to be the most abundant neuronal cell type (Meldrum, 2000; Zhou and Danbolt, 2014). Eight of the seventeen rare clusters (cluster 16, 17, 18, 19, 20, 26, 30 and 32) can be mapped to previously annotated cell types. These include stromal, glutamatergic, macrophage/microglia, radial glia, dopaminergic, granulocyte and GABAergic neuron. Of note, GiniClust3 was able to identify granulocyte cells (cluster 30), even though they represent a tiny fraction (55 out of 1,244,774 cells, 0.004%) of the cell population, indicating the sensitivity of GiniClust3 is very high.

To evaluate the robustness of GiniClust3, we repeated the analysis using randomly subsampled data. To this end, 50% of the cells were randomly selected from common clusters. Since our main focus was to identify rare cell clusters, the cells assigned to these clusters were all retained. Applying GiniClust3 to the subsampled dataset shows that most of the clusters in subsampled dataset are consistent with the original ones (Fig. s1b, s1c). Taken together, these analyses show that GiniClust3 is a sensitive, accurate and efficient clustering method that can be used in many applications.

## Acknowledgements

We thank Dr. Daphne Tsoucas for helpful discussions.

## Funding

This work was supported by a Claudia Barr Award and NIH grant R01HG009663 to GCY.

## Conflict of Interest

none declared.

