## Supplementary figures and images for "GiniClust3: a fast and memory-efficient tool for rare cell type identification"

### Supplementary Figure 1

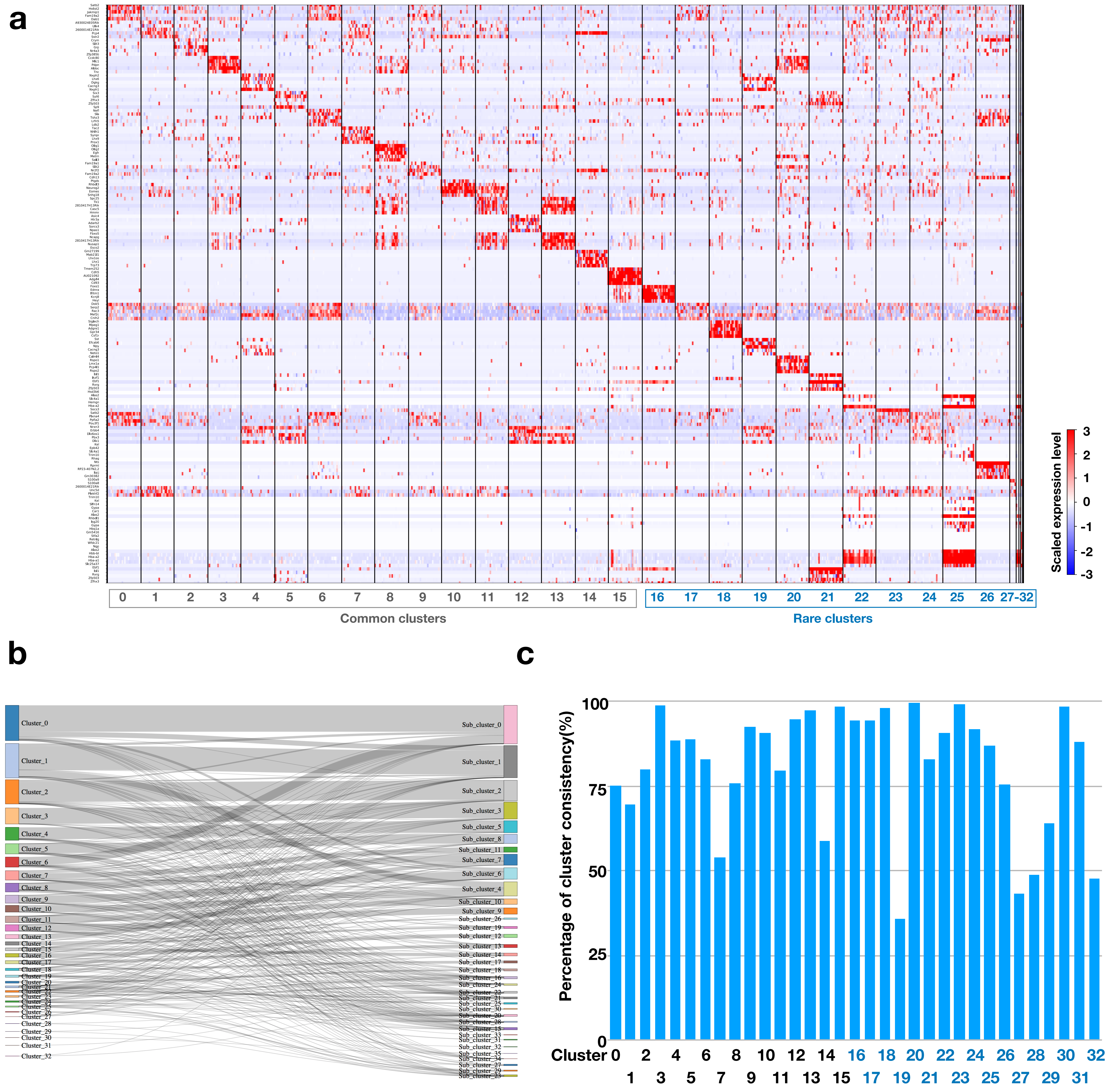
